# Antibody-Fab and -Fc features promote *Mycobacterium tuberculosis* restriction

**DOI:** 10.1101/2024.10.07.617070

**Authors:** Patricia S. Grace, Joshua M. Peters, Jaimie Sixsmith, Richard Lu, Corinne Luedeman, Brooke A. Fenderson, Andrew Vickers, Matthew D. Slein, Edward B. Irvine, Tanya McKitrick, Mo-Hui Wei, Richard D. Cummings, Aaron Wallace, Lisa A. Cavacini, Alok Choudhary, Megan K. Proulx, Christopher Sundling, Gunilla Källenius, Rajko Reljic, Joel D. Ernst, Arturo Casadevall, Camille Locht, Abraham Pinter, Christopher M. Sasseti, Bryan D. Bryson, Sarah M. Fortune, Galit Alter

## Abstract

*Mycobacterium tuberculosis (Mtb),* the causative agent of tuberculosis (TB), is a leading cause of death by an infectious disease globally, with no efficacious vaccine. Antibodies are implicated in *Mtb* control, but the mechanisms of antibody action remain poorly understood. We assembled a library of TB monoclonal antibodies (mAb) and screened for the ability to restrict *Mtb* in mice, identifying protective antibodies targeting known and novel antigens. To dissect the mechanism of mAb-mediated *Mtb* restriction, we optimized a protective lipoarabinomannan-specific mAb through Fc-swapping. *In vivo* analysis of these Fc-variants revealed a critical role for Fc-effector function in *Mtb* restriction. Restrictive Fc-variants altered distribution of *Mtb* across innate immune cells. Single-cell transcriptomics highlighted distinctly activated molecular circuitry within innate immune cell subpopulations, highlighting early activation of neutrophils as a key signature of mAb-mediated *Mtb* restriction. Therefore, improved antibody-mediated restriction of *Mtb* is associated with reorganization of the tissue-level immune response to infection and depends on the collaboration of antibody Fab and Fc.

## Introduction

With an estimated 10 million new cases worldwide and approximately 1.6 million associated deaths annually, tuberculosis (TB) remains one of the world’s deadliest infectious diseases^1^. Novel drugs, host directed therapies, and vaccines are needed to combat this global health threat. While cell-mediated immunity is critical for limiting intracellular *Mycobacterium tuberculosis* (*Mtb*) infection^2^, recent data suggest the importance of humoral responses in mediating control. Humoral immunity has been relatively understudied in TB, but passive transfer of IVIG^3,4^, antibodies from TB patients^5,6^, and vaccine-induced antibodies^7^ in mice demonstrate that antibodies can promote *Mtb* restriction, although these effects vary between studies ^8^. Similarly, monoclonal antibodies (mAb), targeting *Mtb* cell-wall associated antigens lipoarabinomannan (LAM)^9,10^ and heparin-binding hemagglutinin (HBHA)^11^, the membrane-associated PstS1^12^, and intracellular protein HspX^13^ have been shown to improve infection outcomes in mice. However, it remains unclear how antibodies mediate *Mtb* infection control.

Antibody function is governed not only by antigen specificity, conferred by the antigen-binding (Fab) domain to neutralize pathogens, but also by the antibody constant (Fc)-domain, which recruits complement and engages Fc-receptors (FcRs) on the surface of immune cells. These interactions promote Fc-effector functions including phagocytosis, cellular cytotoxic responses in NK cells and macrophage activation^14,15^. The mere presence of *Mtb*-specific antibody titers in a host does not guarantee protection against infection or disease^8^. Instead, emerging data suggest that antibody Fc-effector functions diverge significantly across TB disease states may better predict antibody-mediated protection.^16^ Recently, humoral immune responses have been linked to reduced rates of infection in a TB vaccination study in humans^17^ as well as in non-human primates^18^. Mice lacking the Fc gamma-chain (Fcγ−chain), the FcR moiety key to intracellular signaling and function, exhibit a diminished capacity to restrict *Mtb* compared to wild-type (WT) mice^19^, indicating that IgG interactions with FcR contributes to immune control of infection. Consistent with this, point-mutations of the IgG Fc-domain that reduce FcR engagement of protective mAbs eliminated antibody-mediated *Mtb* restriction^12,20^. Further, *Mtb*-specific Fabs generated with IgA and IgM Fc-domains exhibit increased *Mtb*-restrictive capacity *in vitro* ^18,21^, emphasizing that Fc-effector function is also important for antimicrobial activity of antibodies during *Mtb* infection. However, it is unclear whether antibodies to diverse antigens or even distinct epitopes within the same antigen confer protection, and how Fc-effector functions contribute to *Mtb* restriction *in vivo* is unknown.

To investigate the role of antibody Fab and Fc in immune-mediated *Mtb* restriction, we profiled the largest library of *Mtb*-specific mAbs to date for the ability to drive bacterial control in mice. Screening this library targeting secreted, surface, and lipid antigen revealed diverse and novel antigen targets of mAbs able to restrict *Mtb* growth *in vivo*, that were not predicted by antibody opsonization. We selected a protective LAM-specific mAb and swapped Fc-domains, generating Fc-variants, to interrogate the contribution of Fc-effector function to *Mtb* control. The most functional Fc-variant (mIgG2a) of the αLAM-mAb, defined by combinatorial *in vitro* activities measured in a “Systems Serology” analysis, exhibited enhanced restriction of *Mtb* in mice as compared to the mIgG1 (a less functional Fc-variant). These Fc-variants did not markedly improve macrophage intrinsic *Mtb* control *ex vivo*. Instead, treatment of mice with the mIgG2a Fc-variant prior to infection altered *Mtb* tropism to different innate immune cell populations in the lung. Further single cell transcriptional profiling of lung innate immune cell populations pointed to distinct pathway activation in alveolar macrophage and neutrophil in the presence of the restrictive αLAM-mIgG2a at early stages of *Mtb* infection. These data suggest that protective antibodies cooperate with the innate immune response, rewiring interactions of *Mtb* with innate immune cells within the lung.

## Results

### Diverse antigenic targets of mAbs promote Mtb restriction in vivo

*Mtb* expresses an array of protein and glycolipid antigens that are targeted by antibody responses in humans ^22,23^, and passive transfer of these antibodies to *Mtb*-infected mice suggests that some polyclonal pools harbor protective antigen-specific antibodies^3,5,6,24^. The transfer of mAbs specific for a limited number of antigen targets are described to give rise to various *Mtb* control phenotypes: restricting bacterial replication in the lung, preventing spleen dissemination, or increasing animal survival^9,11–13,25^. Despite this evidence, the identity of the many potential *Mtb*-antigen targets or even epitopes within an individual antigen can promote *Mtb* restriction in the lung remains incompletely characterized. Many studies have focused on the protection mediated by antibodies targeting surface-associated glycan and protein antigens. Whether antibodies targeting intracellular or secreted antigens provide protection equivalent to these surface-associated targets is unclear. Thus, we generated a library of 24 mAbs, targeting various *Mtb* antigens (**Table 1**) to identify novel antigens targeted by antibodies that could promote antimicrobial activity in mice.

**Table 1.** TB monoclonal antibody clones and antigen target. *, Clones publicly available from BEI Resources, **previously unpublished clones characterized in supplemental Table 1, Supplemental Figures 1 and 2.

| mAb Clone | Antigen | Reference |
| --- | --- | --- |
| 712 | Ag85B | (26) |
| 710 | Ag85B | (26) |
| CS-90 | Ag85 complex | * |
| CP2-R19 | ESAT6/CFP10 | ** |
| a-Rv1411c | LprG | * |
| D2 | HBHA | (11) |
| Apa1 30 | Apa |  |
| CP2-LR5 | PstS1 | ** |
| IT-15 | PhoS1/PstS1 | ** |
| Mpt64(B) | Mpt64 | * |
| 2e9 | HspX | (13) |
| OM2-L2 | HspX | ** |
| KatG2 | KatG | * |
| C1-LR/NP12 | Unknown | ** |
| CP2-LR17 | Capsule polysaccharides/lipopolysaccharides | ** |
| 24c5 | alpha-glucan | (10) |
| CP2-R3 | Glycopeptidolipids | ** |
| OM2-L7 | Glycopeptidolipids | ** |
| OM2-L22 | Glycopeptidolipids | ** |
| SMITB14 | LAM (AM) | (12) |
| A194 | LAM | (27) |
| MoAb1 | LAM (MTX) | (27) |
| CS-35 | LAM | * |
| OM2-R18 | Inner membrane and intracellular peptide glycans | ** |

For this study we included mAbs previously described to confer varying protective *Mtb* phenotypes in mice ^9–11,13,26–28^. Additionally, we screened novel mAbs derived from vaccination against capsular (CP) and outer membrane (OM) fractions of *Mtb*. These novel mAbs were characterized for the ability to bind intact *Mtb* cells, *Mtb* lysate proteins in Western Blots, and *Mtb* antigens in Luminex assays **(Table S1 and Fig. S1**). For consistency and comparability, the Fab domain of each mAb was produced with a human IgG1 (hIgG1) Fc-domain, known to interact with FcR and mediate Fc-effector functions in mouse immune cells^29^.

We screened the hIgG1 mAb library to identify mAbs that mediated *Mtb* growth restriction in mice. A 5 mg/kg dose of each mAb was passively transferred to mice one day prior to low-dose aerosol challenge with *Mtb* and we enumerated the bacterial burden in the lung 14 days post *Mtb* challenge. Ten antibodies of the mAb library promoted significant fold reduction of *Mtb* colony forming units (CFU) compared with PBS-treated mice (**Fig. 1A**). Both protein- and glycolipid-specific antibodies restricted *Mtb in vivo*.

**Figure 1.**
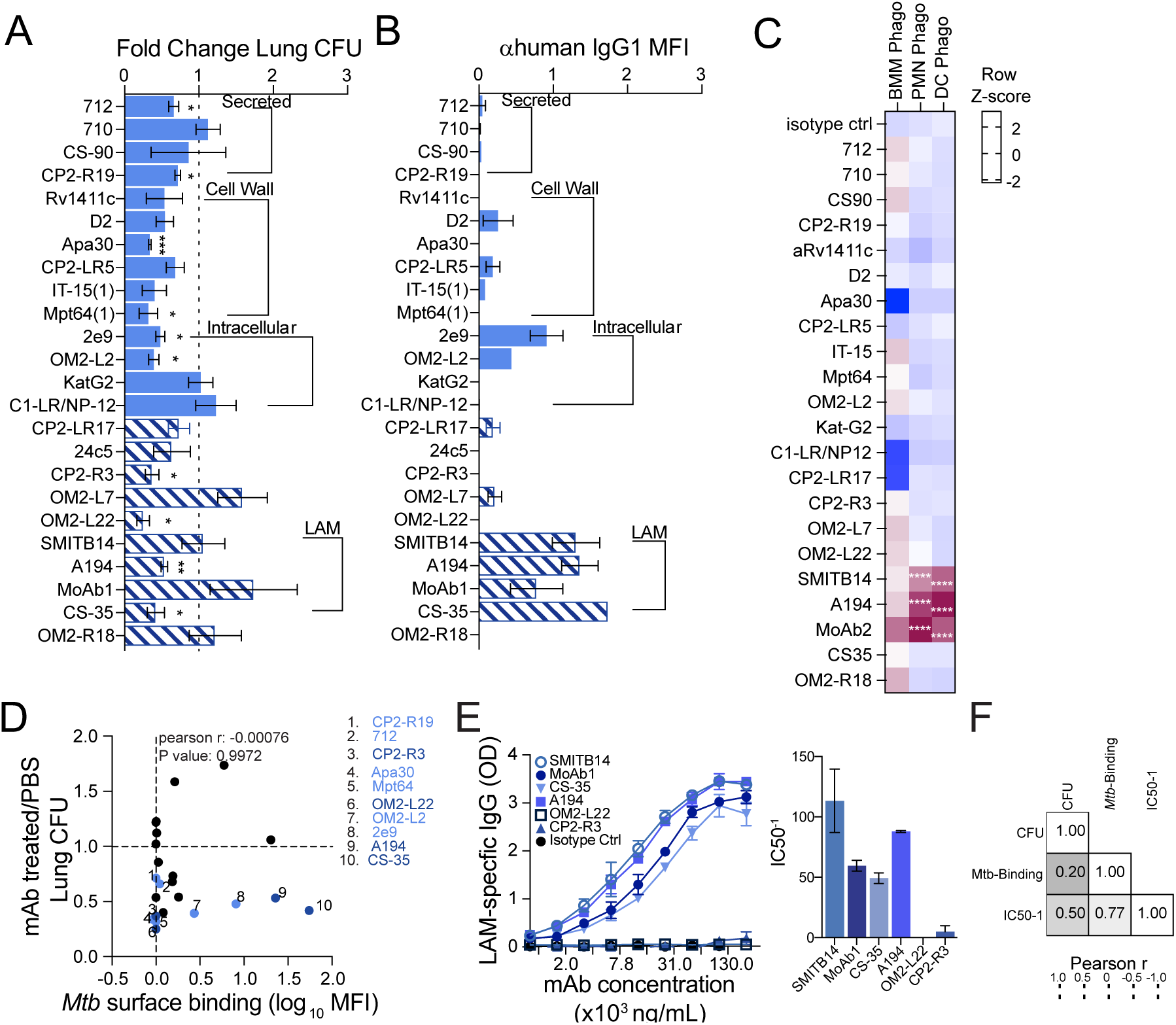
Passive transfer screen of mAb mice identifies protein and glycolipid binding antibodies that restrict *Mtb* outgrowth. A library of hIgG1 monoclonal antibodies targeting proteins (solid light blue bars) and glycolipid antigens (hashed navy bars) were screened for *Mtb* growth restriction in mice. C57BL/6 mice were pre-treated with 100 μg of mAb and infected via aerosol with ∼100 Day 1 CFU of *Mtb* and brackets denote class/localization of TB antigen targets for each mAb clone. *Table 1* defines these antigen targets in more detail. **(A)** Lung Colony Forming Units (CFU) of mAb treated mice normalized to control mice (PBS) in each round of infection (linear scale). Data are representative of 2 independent experiments (n = 3 mice). Graphs depict the mean +/- SEM, and mAb mediating significant restriction were identified by one sample t test: *p, <0.05, **p, <0.01, ***p, <0.005. **(B)** Bar plots represent fold-enrichment of hIgG1 MFI on the surface of *Mtb* measured for each monoclonal via flow cytometry. Each MFI values were normalized to the isotype control stain. **(C)** Fluorescent *Mtb* and mAb were combined prior to exposure to mouse dendritic cells, neutrophils, and macrophages derived bone marrow (Supplemental Figure 2A). Heat map depicts the phagocytosis (phago) scores calculated for each immune cell type, which was z-scored per cell type tested. Phago Score = ((%Mtb^+^cells x Mtb^+^Cell mean fluorescence intensity)/100). Data are representative of 2 independent experiments (n = 2 replicates). mAb clones which significantly enhanced Mtb phagocytosis were identified by One-way ANOVA: ****,p,< 0.0001. **(D)** *Mtb* surface binding plotted against lung CFU are poorly correlated. Light blue dots indicate protein binding mAbs and dark blue dots represent glycolipid specific mAbs that significantly reduced *Mtb* burden in mouse lung. **(E)** Relative monoclonal antibody binding measured by LAM ELISA. The IC50 for titrated LAM-specific mAbs was calculated and the inverse for each antibody describes the binding capacity of each Fab for LAM antigen. **(F)** Pearson correlation r of LAM-binding mAbs with *Mtb*-surface binding and LAM binding IC50.

*Mtb* surface binding was previously associated with *Mtb* restriction *in vivo*^6,30^. Thus, we tested whether restrictive mAbs bound to *Mtb* more effectively than non-restrictive mAbs. A head-to-head comparison of mAb binding was performed with live *Mtb,* revealing, as expected, that LAM-specific mAbs robustly opsonized *Mtb* (**Fig. 1B**, hashed bars), while many of the protein-targeting mAbs did not (solid blue bars). Specifically*, Mtb* opsonization revealed low level binding by mAbs targeting secreted proteins (Ag85B, clone 712) and variable binding was observed for cell-wall- or intracellular-protein specific mAbs (e.g. Apa, clone Apa30 and HspX, clones 2e9 and OM2-L2) (**Fig. 1B).**

Previous studies found a correlation between mAb opsinophagocytic activity and bacterial restriction^30,31^. We tested these mAbs for the ability to increase bacterial phagocytosis in mouse macrophages, neutrophils, and dendritic cells. LAM-specific mAbs promoted significant increases in *Mtb* phagocytosis (phago score) in dendritic cells and neutrophils compared to isotype control treatment, but no significant antibody-mediated increases in *Mtb* uptake were observed in macrophages (**Fig. 1C and Fig S2A**). We found a moderate correlation between *Mtb* surface binding and *Mtb* phagocytosis in dendritic cells and neutrophils, suggesting that antibody opsonization can enhance bacterial uptake by these cells (**Fig. S2B**). However, when comparing across this library there was no significant correlation between phagocytic function and *in vivo* control (**Fig. S2B**). These results indicated that both opsonizing and non-opsonizing antibodies could promote *Mtb* restriction *in vivo*.

Surprisingly, the degree of *Mtb*-surface binding did not correlate with *in vivo* restriction (**Fig. 1D**). Specifically, not all antibodies targeting the surface-abundant LAM antigen restricted *Mtb* outgrowth *in viv*o. Instead, the LAM-specific clones promoted highly variable *Mtb* growth phenotypes from restriction to near enhancing effects in mice. To further dissect the determinants of LAM-specific *in vivo* restriction, we measured the binding affinity of the LAM-specific mAbs for the cognate LAM antigen. We found no significant differences in binding affinity across the LAM-specific mAbs (**Fig. 1D**). While affinity weakly correlated with *Mtb* surface binding, antigen-specific affinity did not correlate with the growth restriction observed in mice (**Fig. 1E**). Collectively, profiling this large mAb library indicates that not only high affinity opsinophagocytic antibodies restrict *Mtb* growth, but that antibodies targeting secreted, cell-wall associated, or canonically intracellular proteins can promote *Mtb* restriction *in vivo*.

### Fc-domain swapping of an αLAM mAb enhances antibody-mediated effector function

Previous data point to a role for antibody cooperation with the innate immune system, via Fc-effector functions, as a key mechanism in the control of *Mtb*. To dissect the impact of Fc-effector function on *Mtb* control *in vivo* we focused on the LAM-specific mAb clone A194^27^, which displayed restrictive efficacy *in vivo* and enhanced phagocytic function in our screen. We reasoned that swapping the Fc-domain of this αLAM mAb for specific mouse isotypes would modulate Fc-effector function and might impact the restrictive effect of the LAM-specific mAb. In addition to the hIgG1 variant that we screened previously, we generated a functional mouse IgG2a (mIgG2a) Fc-variant, an FcR-binding knockout mIgG2a N297A point-mutant, and a mouse IgG1 (mIgG1) Fc-variant. We assayed FcR-binding and antibody effector functions of the αLAM Fc-variants complexed with LAM-coated beads or *Mtb*. Consistent with previous reports^29,32^, the mIgG1 and mIgG2a N297A Fc-variants displayed negligible binding to the activating FcγRIV receptor, while the hIgG1 and mIgG2a variants bound with high affinity (**Fig. 2A**). The mIgG2a, mIgG1, and hIgG1 variants bound with decreasing strength to FcγRIIIA, while the mIgG2a N297A displayed no apparent binding (**Fig. 2B**). Finally, the mIgG1 variant bound with the greatest affinity to FcγRIIB; the mIgG2a and hIgG1 bound with relatively moderate affinity, while the mIgG2a N297A had no apparent binding to FcγRIIB (**Fig. 2C**).

**Figure 2:**
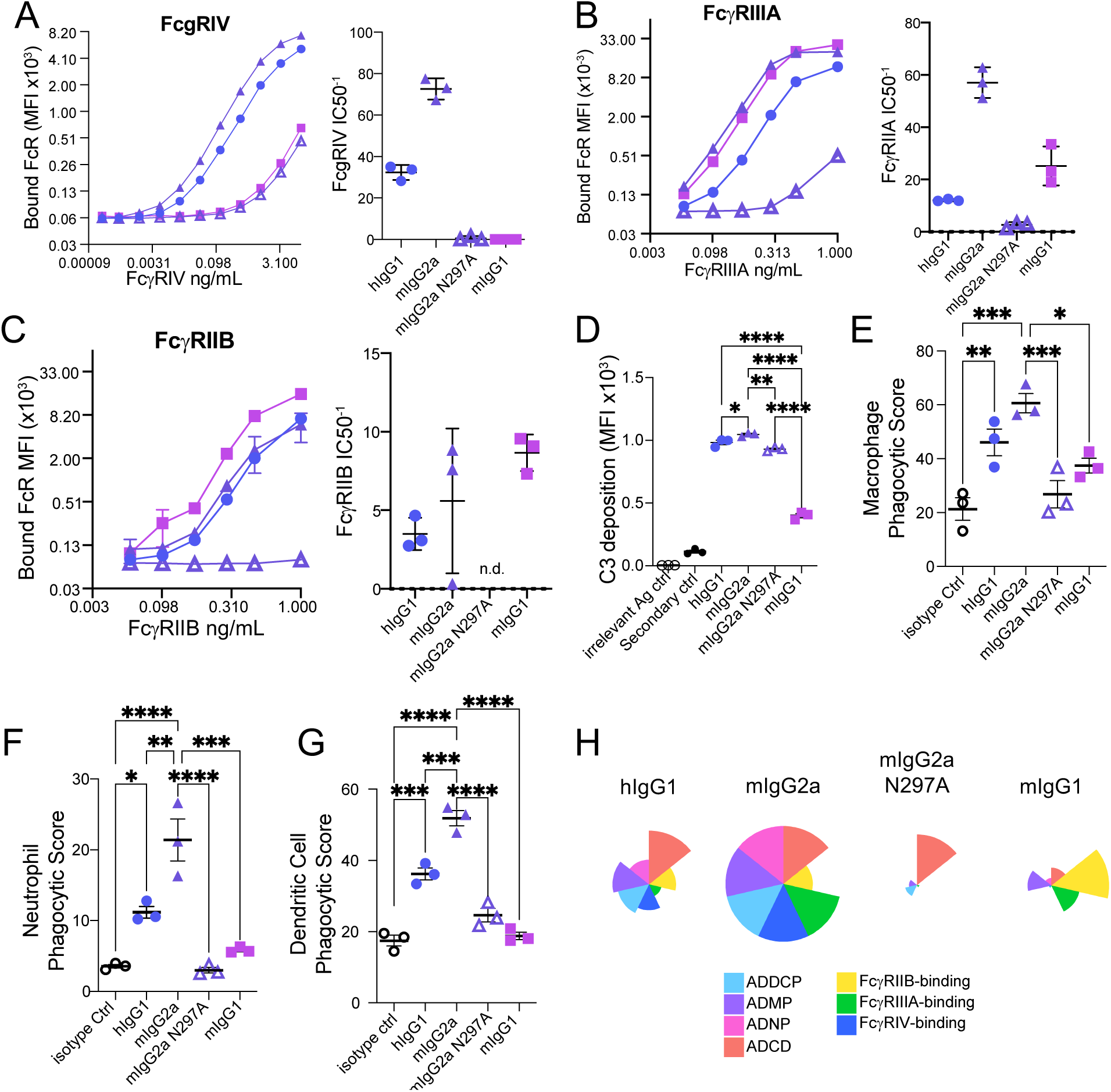
FcR binding and Fc-effector functions of αLAM Fc-variants. Luminex beads were combined with Fc-variant mAbs to test for the ability to bind mouse FcRs. Graphs depict MFI of **(A)** PE-FcγRIV, **(B)** -FcγRIIIA, and **(C)** -FcγRIIB bound to LAM beads complexed with αLAM Fc-variants. **(D)** Relative MFI of αC3-FITC antibody used to detect C3 deposition on LAM-bead immune complexes (IC). Phagocytic Scores determined for αLAM Fc-variant opsonized *Mtb* in bone marrow derived **(E)** macrophages, **(F)** neutrophils, and **(G)** dendritic cells. Data are representative of 2 independent experiments. Graphs depict the mean +/- SEM. Significant differences between Fc-variants were determined by one-way ANOVA with Tukey’s multiple correction: *p, 0.05, **p, 0.01, ***p, 0.005.

Differences in FcR affinity impart differential Fc-effector function^29^. In addition to biophysical measures of FcR interactions, we compared the capacity of Fc-variants to mediate complement recruitment and *Mtb*-phagocytic functions. When the mAbs were complexed with LAM-coated beads, all Fc-variants were able to recruit C3 complement on immune complexes, indicating mature complement-complex formation, but the mIgG1 variant displayed a lower level of C3 recruitment (**Fig. 2D**). Notably, while the mIgG2a N297A displayed reduced FcR interactions, the ability of the variant to recruit complement was maintained. We also screened Fc-mediated phagocytic function in murine macrophages, neutrophils, and dendritic cells and found that the mIgG2a-variant possessed enhanced phagocytic activity in macrophages (**Fig. 2E**), neutrophils (**Fig. 2F**), and dendritic cells when compared to the hIgG1 variant (**Fig. 2G**). As expected, the FcR-binding mutant mIgG2a N297A, had no appreciable phagocytic capacity compared to an isotype control antibody in any of the cell types tested. The mIgG1 variant displayed modest phagocytic capacity when compared with the mIgG2a Fc-variant. Across the biophysical and functional measures, we confirmed that swapping the Fc domains altered the functional capacity of the αLAM mAb, with each variant displaying distinct FcR and cellular engagement patterns (**Fig. 2H**).

### αLAM-Fc-variants leverage distinct cellular immune responses in vivo during Mtb infection

The diverse functional profiles of the Fc-variants provided an opportunity to probe the role of Fc−effector function in modulating *Mtb* infection *in vivo*. We selected the enhanced (mIgG2a), intermediate (mIgG1), and reduced (mIgG2a N297A) Fc-variants to investigate Fc-mediated immune responses to *Mtb* infection. Fc-variants were passively transferred into mice and animals were challenged with low-dose aerosolized *Mtb.* Following infection, we tracked how immune responses changed across the distinct Fc-variant-treated groups. Using multi-parameter flow cytometry we tracked immune cell recruitment and fluorescent *Mtb* distribution amongst alveolar macrophages (AMs), eosinophils, polymorphonuclear neutrophils (PMNs), classical Ly6C^high^ monocytes, non-classical monocytes (Ly6C^low^,FcgRIV^high^)^33^, interstitial macrophages (IM), CD11b^+^ dendritic cells (CD11b^+^DC), and CD103^+^ DC recruited to the lung (**Fig. S3)** ^34,35^. Analysis of lung cells 14 days post *Mtb* challenge revealed a shift in relative proportions of immune cells in the setting of specific Fc-variant treatments compared to controls (**Fig. 3A**), although the total number of myeloid cells was not statistically different across the treatment groups (**Fig. 3B**). Specifically, we observed elevated AM numbers within the lungs of mice treated with mIgG2a and mIgG2a N297A Fc-variants compared to mice treated with mIgG1 or control mice (**Fig. 3C**).

**Figure 3.**
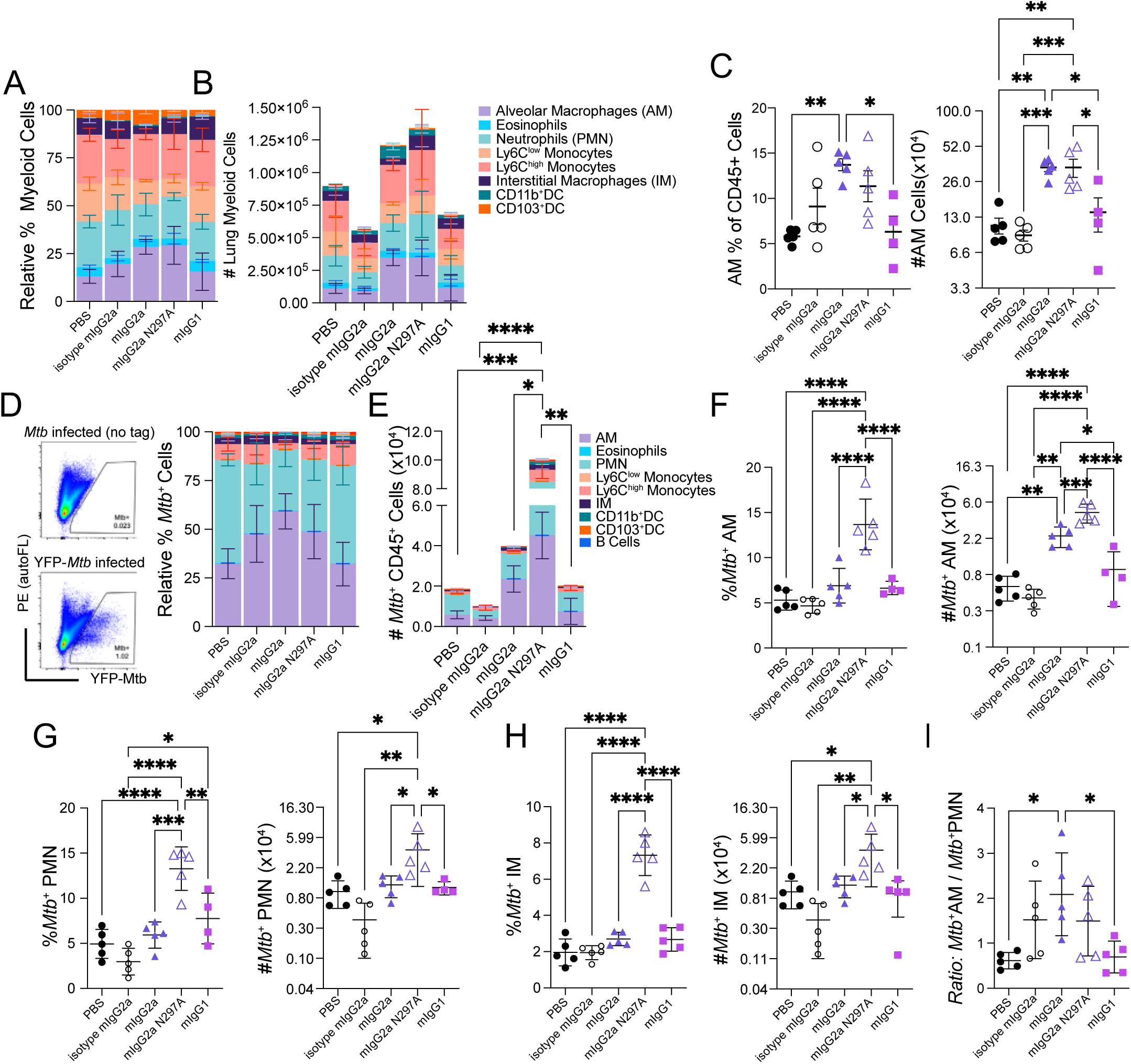
*Mtb* infection of Alveolar Macrophages is enriched within αLAM mIgG2a treated mice. C57BL/6 mice were pre-treated with 100 μg of mAb A914 and infected via aerosol with ∼100 Day 1 CFU of *Mtb.* Single cell suspensions of lung tissue from mAb-treated mice were generated and phenotypically analyzed using flow cytometry (Fig. S3). Stacked bar plots show **(A)** relative frequency (%) of myeloid cell types and **(B)** total numbers (#) of myeloid cells by type present in the lung 14 days post infection. **(C)** Frequency (%) and number (#) of alveolar macrophages (AM). **(D)** Flow plots depict YFP-*Mtb* signal detected in lung cells at 14 days and bar plots show relative frequency of *Mtb*-infected cells by type. **(E)** Total numbers of *Mtb*- infected cells by type. **(F)** % and # of *Mtb*^+^ AM. **(G)** % and # of *Mtb*^+^PMN. **(H)** % and # of *Mtb+* IM. **(I)** Ratio of *Mtb*^+^AM to *Mtb*^+^PMN. Data are representative of 2 independent experiments (n = 4-5 mice per treatment group). Graphs depict the mean +/- SEM, and significant differences between Fc-variants were determined by one-way ANOVA with Tukey’s multiple correction: *p, 0.05, **p, 0.01, ***p, 0.005, ****p, 0.0001.

Using yellow fluorescent protein expressing (YFP)-*Mtb,* we also tracked *Mtb-*distribution across lung immune cells. Consistent with previous reports in C57BL/6 mice 14 days post infection^36,37^, YFP-*Mtb* was found within AM, PMN, and IM of all animals regardless of treatment *Mtb* (**Fig. 3D**). However, the Fc-variants differentially affected the relative distribution of *Mtb* amongst the three cell types. The most striking effects were associated with mIgG2a-N297A treatment which resulted in significantly higher numbers of *Mtb*+ immune cells (Mtb^+^CD45^+^) (**Fig. 3E)** and greater relative numbers of *Mtb*+ AM (**Fig. 3F**), PMN (**Fig. 3G**) and RM (**Fig. 3H**) compared to treatment with any other Fc-variants. These data suggest that loss of FcR engagement but retained complement recruitment activity (**Fig. 2D**) may promote differential bacterial entry into immune cells. The effects of the other Fc-variants were less pronounced, although mIgG2a treatment resulted in greater numbers of *Mtb+* alveolar macrophages (**Fig. 3F)** and higher ratios of infected AM to PMN when compared with other Fc-variant treatments (**Fig. 3I**). These results indicated that mAbs with enhanced Fc-effector function promotes residence of *Mtb* within AM and may either limit *Mtb*-uptake by or promote bacterial growth restriction within neutrophils.

### αLAM mIgG2a treatment is associated with temporal bacterial restriction in vivo

We next determined the effect of the different αLAM Fc-variants on bacterial restriction in the lungs of mice following *Mtb* aerosol challenge (**Fig. 4A**). Treatment with the αLAM-mIgG2a Fc-variants, like the hIgG1 screened previously (**Fig 1A**), resulted in significantly lower bacterial burden in the lung when compared to control mice 14 days post infection (**Fig. 4B**). Conversely, no restriction was observed with mIgG2a-N297A, the Fc-variant with reduced FcR-binding, or the less functional mIgG1 Fc-variant, indicating that Fc-effector function is required for αLAM- mediated bacterial control. *Mtb* dissemination from the lung to other tissues occur in a non-linear fashion 11-17-days following infection^38^; notably, at the early time point of 14 days, we detected *Mtb* only within spleens of mIgG2a N297A treated mice (**Fig. 4C**). These data indicated that Fc-effector function is key to the αLAM-mediated restrictive effect on *Mtb*.

**Figure 4.**
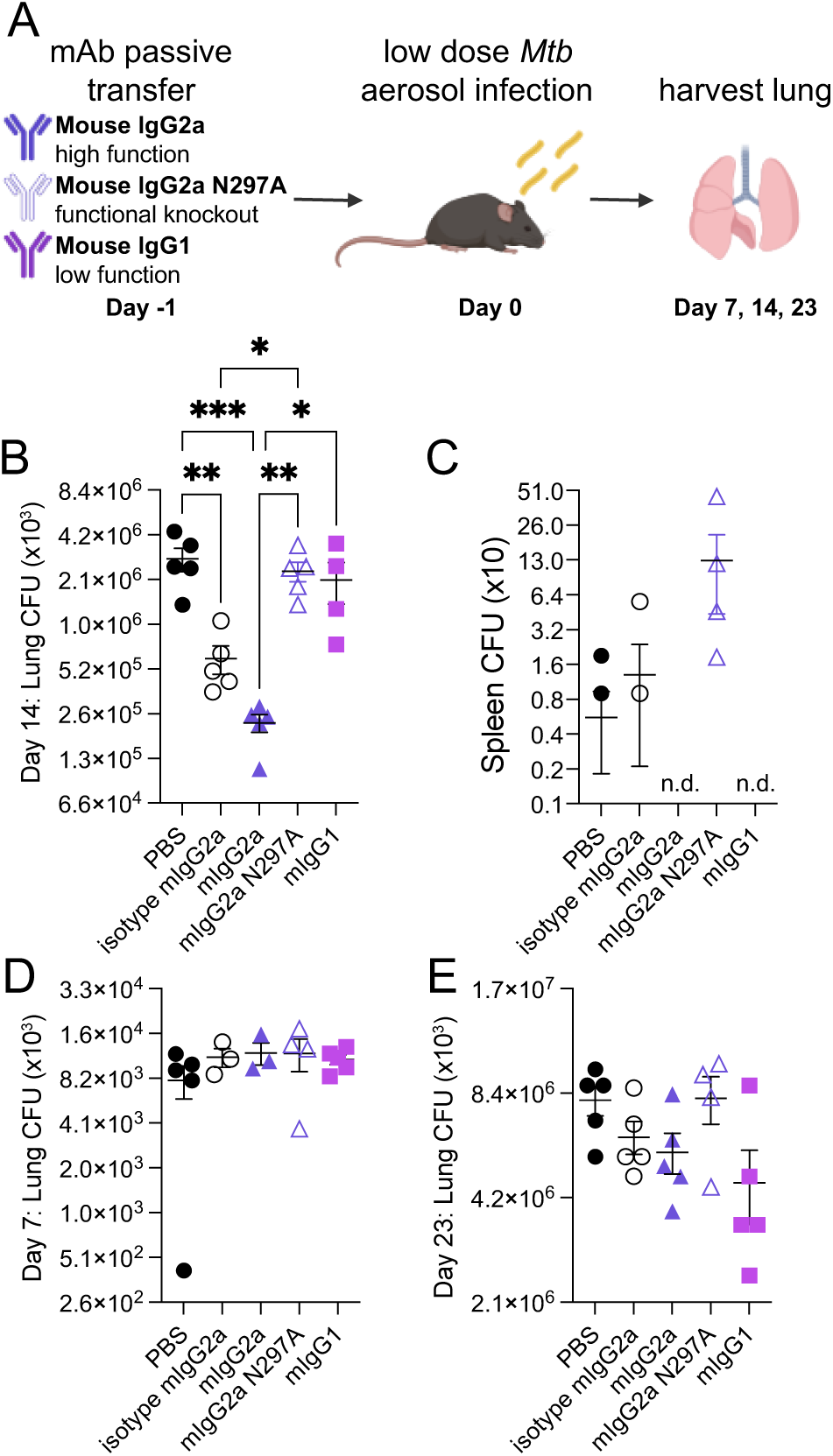
αLAM-mIgG2a restricts *Mtb growth* in early infection. **(A)** C57BL/6 mice were pre-treated with 100 μg of mAb A914 and infected via aerosol with ∼100 Day 1 CFU of *Mtb.* CFU 14 days post infection with *Mtb* in the **(B)** Lung and **(C)** Spleen in αLAM Fc-variant treated mice. **(D)** Lung CFU 7 days post infection. **(E)** Lung CFU 23 days post infection. Data are representative of 2 independent experiments (n = 4-5 mice per treatment group) Graphs depict the mean +/- SEM, and significant differences between Fc-variants were determined by one-way ANOVA with Tukey’s multiple correction: *p, 0.05, **p, 0.01, ***p, 0.005, ****p, 0.0001.

The effect of Fc-variant treatment was changed over time. While the effect on *Mtb* restriction was evident 14 days post infection, there were no differences in lung CFU across the treatment groups within the first week of infection (**Fig 4D**). Moreover, the effect of Fc-variant treatment on *Mtb* growth was no longer significant by 3 weeks post infection (**Fig. 4E**), when passively-transferred antibodies are reported to wane from circulation^39,40^. Collectively, these data are consistent with a model where the presence of antibodies capable of inducing strong Fc-effector functions can promote *Mtb* restriction during the early stages of infection.

### Fc-variants mediate no appreciable Mtb growth difference in in vitro cell infection

Given the differences in bacterial outgrowth and the differential distribution of *Mtb* amongst immune cells we observed in Fc-variant treated mice, we measured the *in vitro*-effect of Fc-variants on antibody-mediated *Mtb* growth restriction in specific lung immune cells. We focused on macrophage and neutrophils, which were the dominantly infected cell types in mouse lungs. We sorted AM, IM, and PMN from the lungs of naïve mice. We combined Fc-variants and luciferase-expressing *Mtb* (lux-*Mtb*), forming Fc-variant immune complexes, which we exposed to the sorted immune cells. We tracked *Mtb* outgrowth via luminescence over time and observed no Fc-variant-mediated restriction of *Mtb* in the sorted AM (**Fig. S4A**), IM (**Fig S4B**), and PMN (**Fig. S4C**). The lack of mAb-mediated *Mtb* restriction *in vitro* indicated a potential non-canonical mechanism of mAb-mediated bacterial restriction that was not observable using individually cultured cells.

### Fc-dependent signaling in AM and neutrophil populations early in Mtb infection

To evaluate how Fc-function may shape immune signaling *in vivo and* bacterial control, we performed single cell RNA sequencing (scRNAseq) on lung immune cells from *Mtb*-infected mice treated with the restrictive αLAM mIgG2a-variant, non-restrictive mIgG2a N297A-variant, or PBS. We focused on cell state early during infection (5 days) to identify transcriptomic differences that are likely to contribute to, rather than result from, CFU differences observed 14 days post infection (**Fig. 5A**).

**Figure 5.**
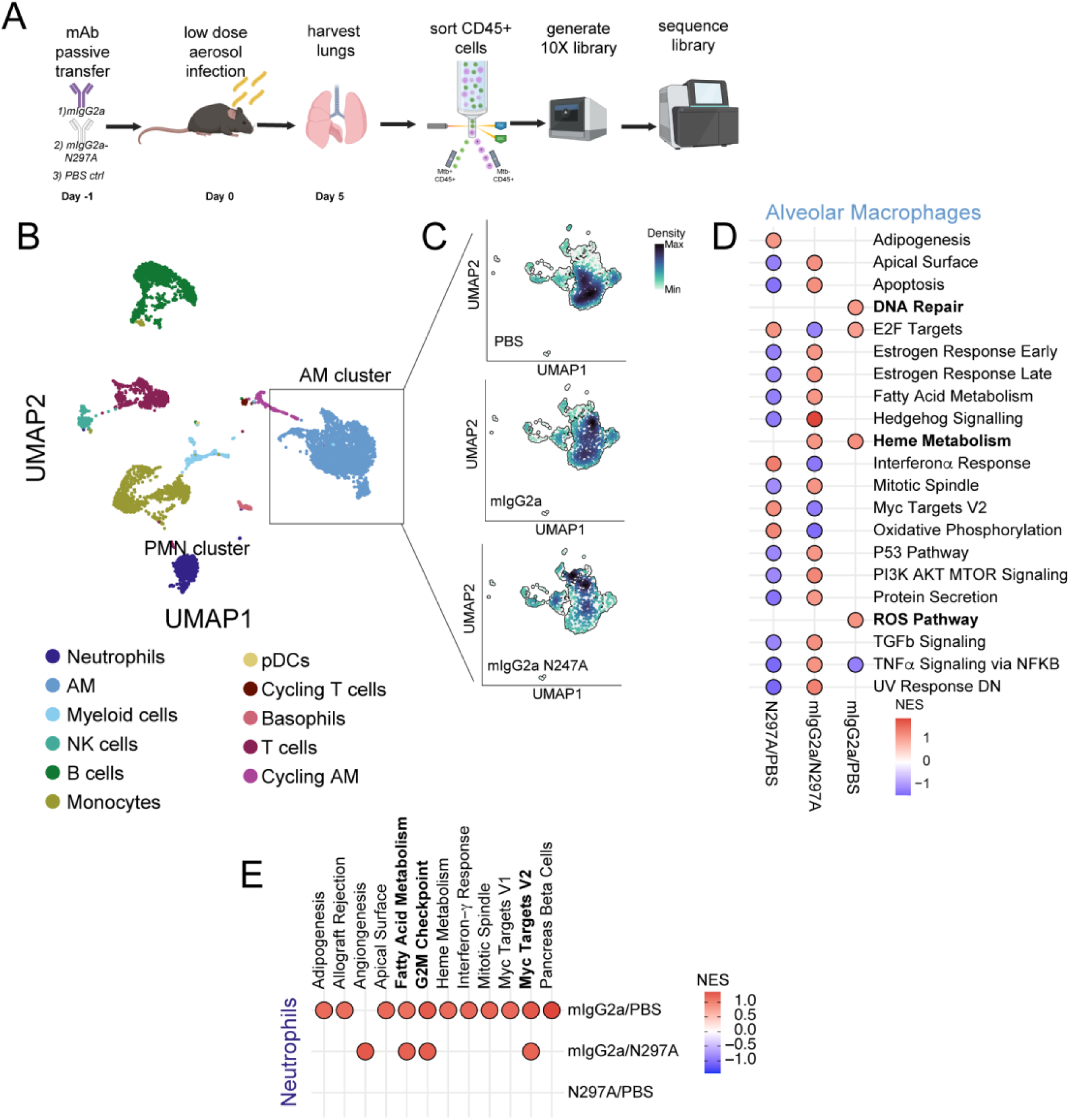
Single cell RNA sequencing of aLAM-mIgG2a and –mIgG2a N297A treated lung cells reveals distinct AM and PMN states. **(A)** C57BL/6 mice were pre-treated with 100mg of mAb A914 and infected via aerosol with ∼60 Day 1 CFU of YFP-*Mtb.* **(B)** UMAP depicts clusters of immune cells defined from single cell RNA-seq: including Neutrophils, alveolar macrophages, myeloid cells, NK cells, B cells, Monocytes, plasmacytoid dendritic cells (pDCS), cycling T cells, Basophils, T cells, and Cycling AM. **(C)** Cells from the distinct Fc-variant treatment groups are differentially distributed within the AM subcluster. **(D)** Top pathways of AM subcluster identified in Gene Set Enrichment Analysis (GSEA) in at least one of the following pairwise comparisons: mIgG2a/PBS, mIgG2a/N297A, N297A/PBS. **(E)** Differentially enriched pathways within neutrophils identified in GSEA in at least one of the following pairwise comparisons: mIgG2a/PBS, mIgG2a/N297A, N297A/PBS. Enriched pathways depicted had normalized enrichment scores (NES)>1.1, signifying enrichment in the numerator, (red dots) or NES<-1.3 signifying enrichment in the denominator (blue dots). Enriched pathways with nominal p values <0.05 are filled with red, significant numerator enrichment or blue for denominator enrichment for the pairwise comparison.

As observed one week post infection (**Fig. 4D**), *Mtb* burden in the lungs were similar in all treated animals five days post infection (**Fig. S5A**). We sorted Mtb+ and Mtb-CD45^+^ cells from PBS, αLAM-mIgG2a, and -mIgG2a N297A treated mice (n=3 mice per treatment) for scRNAseq, recovering 6,702 CD45+ cells from all mice in all conditions. Eleven distinct cell populations were identified by transcriptomic clustering^41^, including neutrophils, AM, myeloid cells, NK Cells, B Cells, Monocytes, pDCs, cycling T Cells, Basophils, T cells, and cycling AM (**Fig. 5B)**, and the relative abundance of the populations recovered by scRNAseq were also observed by flow analysis in the same tissue **(Fig. S5B-C).**

Mapping the distribution of the AM isolated from each of mAb-treated groups, we found distinct antibody-specific enrichment patterns across the subclustered AM (**Fig. 5C**), indicating that antibodies promoted shifts in AM transcriptional states. To identify distinct pathways induced in AMs of mAb-treated mice, we performed gene set enrichment analysis (GSEA) of the expressed genes measured within the pseudo-bulked AM subcluster. We performed pairwise comparisons of cells from mIgG2a-, mIgG2a-N297A-, PBS control-treated and infected mice. In AMs from restrictive mIgG2a antibody treated mice, we consistently observed activation of DNA repair, heme metabolism, and reactive oxygen species (ROS) pathways (red dots) (**Fig. 5D**). In the AMs from non-restrictive mIgG2a N297A-treated mice we observed an enriched gene expression profile which indicated an enhanced IFNα response accompanied by oxidative phosphorylation and Myc-target gene activation (blue dots in mIgG2a/N297A comparisons and red dots in N297A/PBS comparisons) (**Fig. 5D**). The GSEA of AM revealed transcriptional signature shifts, associated with the functional Fc-variant, occurred prior to the onset of *Mtb* restriction in mice.

GSEA analysis on the pseudo-bulked neutrophils revealed additional Fc-mediated changes to lung neutrophil state. Several pathways were enriched within the cells from mIgG2a-treated mice versus mIgG2a N297A-treated or PBS control mice. No pathways reached significance in the comparison of N297A and PBS (**Fig. 5E)**. In the neutrophils of mIgG2a-treated mice, there was differential expression of fatty acid metabolism, G2M checkpoint, and Myc target genes (**Fig. 5E)**. Thus, single cell sequencing revealed early shifts in cellular transcriptional circuitry ahead of observed *in vivo Mtb* control, coordinated by mAb-mediated cellular activation. Activation of these non- canonical pathways in AM and PMN may render cells of the lung more resistant to bacterial growth, resulting in acute *Mtb* restriction and delayed dissemination in mice.

## Discussion

Emerging data indicates that antibody function is a correlate of protection against TB^16,42^. *Mtb*- specific antibodies can enhance *Mtb* restriction *in vitro,* in macrophages^6,16,30^ and human whole blood^43^, and *in vivo,* in mouse models of infection^5,6,12^. While antibodies able to opsonize *Mtb* are commonly associated with antimicrobial activity at early stages of *Mtb* infection *in vivo*^6,7^, the potential for antibodies targeting diverse *Mtb* antigens, including non-surface antigens, remains incompletely understood. By screening a library of mAbs targeting various *Mtb* intracellular, cell wall-associated, and secreted antigens we found that antibody opsonization of *Mtb* and opsinophagocytic activity were concordant, but did not correlate with *in vivo* restriction in mice. Instead, both opsonizing and non-opsonizing mAbs restricted *Mtb*, indicating that additional functions of antibodies may be responsible for antimicrobial activity *in vivo*. Further dissection of the *in vivo* immunological responses induced by a LAM-specific mAb pointed to a clear Fc- dependent mechanism of protection, linked to an Fc-dependent shift of bacterial tropism across lung immune cells. In addition, we observed non-canonical transcriptional activation of immune cells early following *Mtb* infection. Together, these data point to a previously unappreciated role for antibodies in early *Mtb* restriction within the lung, via transcriptional rewiring of key innate immune cell types. In the setting of this LAM-specific mAb, the mAb alone did not clear the bacteria, but collaborates with cells responding to infection in the lung to promote bacterial clearance.

We identified several mAbs, capable of restricting *Mtb in vivo,* despite poor or negligible *Mtb* surface-binding. These mAbs target counterintuitive antigens described as secreted or intracellular: Ag85B, Apa, Mpt64, and HspX. Consistent with our observations, antibodies against HspX and Ag85 were previously shown to protect against *Mtb* during passive transfer experiments in mice^7,13^. During infection, antigen targets have been identified in endocytic vesicles, which transport *Mtb* antigens to the extracellular space^44^, making these targets potentially accessible to antibodies independent of the bacterial surface. Antibody interactions with released antigens may form complexes that co-ligate Fc-receptors on infected cells to promote eradication of intracellular *Mtb* or engage FcRs to promote inflammatory responses^45^. In addition, Mpt64 has previously been identified in infected cell membranes^46^; antibodies specific for this and other cell surface-associated antigens could recruit cells to recognize and eliminate infected cells. In other infectious diseases, such as malaria and HIV, antibodies that bind the surface of infected cells can recruit FcR bearing NK cells to induce antibody-dependent cellular cytotoxicity (ADCC) and enhance infection control^47–49^. Thus, antibodies targeting diversely localized *Mtb* antigens, such as those identified in our screen, may induce Fc-effector functions beyond bacterial phagocytosis and contribute to immune-mediated protection during *Mtb* infection.

Previous studies found an enrichment of antibodies with enhanced binding to FcγRIIIA in patients that controlled TB infection^16^. FcγRIIIA is an activating FcR that directs ADCC through NK cells and macrophages. Phenotypic analyses of peripheral blood mononuclear cells in a cohort of TB patients also identified increased expression of FcγRIIIA on NK cells in humans with controlled *Mtb* infection^50^, further implicating engagement of specific FcRs in TB control. Here, we demonstrate that αLAM Fc-variants with the highest affinity for FcγRIV, the mouse FcR analog to human FcγRIIIA^29,32^, promoted robust restriction of *Mtb* infection *in vivo*. While mouse NK cells do not express FcγRIV, many myeloid cells, including AMs, express FcγRIV (**Fig. S6**). As the first phagocytes to encounter *Mtb* during infection, AMs may respond rapidly and robustly to antibody-opsonized *Mtb* and contribute to rapid bacterial capture or growth restriction^36,51,52^. Importantly, antibody-mediated FcγRIV-ligation on AMs may play a critical role in antibody-mediated protection against Influenza, as the depletion of AMs prior to viral challenge disrupted antibody-mediated restriction of influenza^53^. These data point to a critical role for early antibody-mediated activation of AMs as a first line of defense against *Mtb* and other respiratory pathogens, potentially linked to FcγRIV in mice or FcγRIIIA in humans.

Transcriptional profiling of the lung cellular immune response to mAb Fc-variant treatment following *Mtb* infection revealed distinct shifts in cell state following mAb variant treatment. We found early unique activation profiles associated with the restrictive αLAM mIgG2a compared to controls in AMs and PMNs within the infected tissue. Consistent with previous studies of macrophages^54^, we found ROS signaling elevated in restrictive mIgG2a-treated lung AMs. In a model of lupus nephritis, antibody-Fc has recently been found to shift liver macrophage metabolism following antibody treatment, and this metabolic shift promoted greater inflammatory responses to immune complexes^55^. Here we found a heme metabolism pathway preferentially elevated in AMs in mice treated with the restrictive mIgG2a variant. Similarly, this pathway has been found to be enriched in *Mtb* restrictive AMs in concomitantly immune animal models ^56^ and transcriptional changes in BCG-trained immunity models also reveal heme metabolism enrichment^57^.

PMNs also displayed unique shifts in metabolism. Both fatty acid and heme metabolism gene sets were enriched in the lung PMN of protective mIgG2a-treated mice, indicating that metabolic shifts in neutrophils may promote a less hospitable environment for bacterial growth. Although there is a paucity of data linking metabolism in PMNs to *Mtb* control^58,59^, recent studies have found that distinct metabolic activation of neutrophils can promote inflammation or prevent tissue damage^60^. While this study highlights a relationship of metabolism and antibody Fc recognition on immune cells, future studies are essential to understand how metabolic signals impact the cellular response during *Mtb* infection.

Recent TB vaccine studies point to a role for antibodies in *Mtb* control^17^. Likewise, intravenous BCG vaccination in non-human primates provide near sterilizing protection against *Mtb* and is linked to both cellular^61^ and humoral immune responses^18^. Our current work contributes to the growing appreciation that humoral immunity contributes to protection against TB, extending our understanding of antigen-targets and functional diversity of antibodies that promote bacterial control. Furthermore, the data presented here point to a critical collaboration between the humoral and innate immune response in early defense against *Mtb*. Thus, next generation TB vaccines that exploit canonical (opsinophagocytosis and ADCC) and non-canonical (metabolic) functions may enhance clearance of this deadly pathogen. Here we focused on the role of antibodies as a prophylactic measure against *Mtb* infection. In our model, antibodies engage innate immune cells in the absence of T cells. However, in vaccination or therapy, antibodies would persist for longer periods of time, tonically interacting with the innate immune system, and likely would further collaborate with T cells. Several *Mtb* infection models indicate that antibody-mediated function can be dependent on T cells^3,5^, and future studies will be essential to fully dissect the precise antibody Fc biology that best cooperates with T cell immunity to gain optimal control over *Mtb* infection.

## Materials and Methods

### Antibody cloning and expression

We Fc-engineered previously published antibody clones A914^27^), D2^11^), 710 and 712^26^), MoAb1^27^, 2e9^13^, SMITB14^9^, and 24c5^62^). Additional VH/VL sequences were courtesy of and MassBiologics and Chris Sassetti (OM- and CP- clones), AERAS (Apa30), and BEI Resources (CS-90, a-Rv1411c, IT-15, Mpt64(B), KatG2, CS-35). Antibody VH/VL sequences were cloned together with hIgG1, mIg2a, mIgG2a N297A, and mIgG1 Fc sequences, and the corresponding antibodies were produced following transfection of plasmids into Expi293F cells (in the Dana Farber Cancer Institute Antibody Production Core). Antibodies were enriched from culture supernatants using protein G beads and were tested for endotoxin. Only antibody preparations with less than 0.5 endotoxin unit/mL detected were used in our experiments.

### Bacterial Strains

YFP-expressing *M. tuberculosis* (courtesy of C. Sassetti) and the parental strain H37Rv were used for mouse aerosol infections. For *in vitro* infection of phagocytic cells YFP-*Mtb* was used to track bacterial uptake and *Mtb-276,* a luminescent *Mtb* strain (*lux*-*Mtb,* S. Fortune) was used to track bacterial growth in the *in vitro Mtb* restriction assays.

### Mtb surface staining

10^7^ YFP-*Mtb* cultured in 7H9 with and/or without 0.05% Tween-80, were combined with 1μg of mAb in a 96-well plate at 37°C for one hour. Following incubation with antibody, *Mtb* was washed 2x with PBS and stained with an αhuman IgG1-Fc antibody (M1310G05, Biolegend) secondary antibody. After staining with secondary antibody, *Mtb* was washed with PBS and fixed overnight with 4% paraformaldehyde (PFA) before analysis on a BD LSR Fortessa and High Throughput Screening (HTS) plate reader. Data were analyzed using FlowJo software version 10.8.1 for Mac OS X.

### Mice

C57BL/6 mice used in these studies were purchased from The Jackson Laboratory (Bar Harbor, ME). All animal experiments were conducted in accordance with procedures approved by the Institutional Animal Care and Use Committees of Harvard T.H. Chan School of Public Health and the Ragon Institute of MGH. MIT, and Harvard.

### Aerosol infections

7-week-old female C57BL/6 mice were injected with 100 μg of antibody (5 mg/kg) i.p. one day prior to aerosol infection with YFP-H37Rv. Prior to infection H37Rv was cultured in 7H9-OADC media to mid-log phase and passaged once. Mice were inoculated with 50-200 CFU using an Inhalation Exposure Unit (Glas-Col). 3 animals from the aerosol infection were sacrificed one day following infection to determine the Day 1 CFU dose. 14 days following aerosol infection, all mice were sacrificed, and lungs and spleen were harvested for CFU plating and single cell suspensions were stained for flow cytometric analysis.

### LAM coated bead phagocytic assays

Biotinylated LAM was generated by combining 100 μg of LAM (BEI Resources NR-14848) dissolved ddH_2_O (1mg/mL), with 10 μL of 1M sodium acetate (NaOAc, Millipore Sigma) and 2.2 μL of 50mM sodium periodate (NaIO_4_, Millipore Sigma); this solution was incubated for 60 minutes (mins) on ice in the dark to oxidize LAM. To stop the oxidation reaction, 12uL of 0.8M NaIO4 was added to the solution and incubated for 5mins at room temperature (RT) in the dark. The oxidized LAM was transferred to a fresh tube and combined with 10 μL 1M NaOAc and 22 μL of 50mM hydrazide biotin (Millipore Sigma), this was incubated at RT for 2 hours (hrs) to biotinylate LAM. The reaction mixture was buffer exchanged on an Amicon 3KDa cutoff 0.5mL Centrifuge column (Millipore Sigma) to remove excess biotin by washing 3 times with PBS. The buffer-exchanged biotinylated LAM was suspended in a final volume of 100 μL of PBS. As described^18^, biotinylated LAM was combined with 1 μm green fluorescent neutravidin beads (ThermoFisher) incubated overnight and washed to generate LAM-coated beads. These beads were co-incubated with 0.5 μg of αLAM Fc-variants for 1hr at 37°C to generate immune-complexes, washed with PBS then combined with either RAW cells for antibody-dependent phagocytosis (ADCP) assays or bone marrow derived neutrophils (generated as described below) for antibody-dependent neutrophil phagocytic (ADNP) assays. Cells and immune complexes were co-incubated for 1hr at 37°C, after which the cells were washed with PBS, stained with fixable live/dead NearIR stain (ThermoFisher), and fixed with 4%PFA. Phagocytosis data was collected on a BD LSR Fortessa and HTS plate reader. Data were analyzed using FlowJo software version 10.8.1 for Mac OS X. Phagocytic Scores were determined as (%Green *LAM-bead*^+^Live Cells x MFI of *Bead*^+^ Cells)/100 for each of the antibody conditions tested.

### Mtb Phagocytic Assays in bone marrow derived cells

Bone marrow was isolated from female C57BL/6 mice, aged 6-8 weeks. BMDC and neutrophils were generated in 7-day cultures with complete RPMI-10 supplemented with 15ng/mL recombinant GM-CSF (PeproTech). Cells from the floating fraction were harvested and BMDC were sorted using anti-CD11c MicroBeads (Miltenyi Biotech) or anti-Ly6G MicroBeads (Miltenyi Biotech) for the isolation of BMDC and PMN, respectively. BMM were differentiated in complete DMEM-10 supplemented with 15 ng/mL recombinant M-CSF(PeproTech) and 7 days later adherent macrophages were collected from the culture dish with warm PBS. Differentiated cells were seeded into 96-well plates at 3×10^4^/well. 1.5×10^5^ antibody-coated *Mtb* (prepared as described above in *Mtb surface staining*) were added to each well. Cells and bacteria were co-incubated for phagocytosis for 1hr, after which extracellular bacteria was washed off with PBS and the cells were stained with CD11b and Live/Dead for identification of live cells via flow cytometry. Cells were fixed at 4°C overnight with 4% PFA and run on a BD LSR Fortessa and HTS plate reader. Data were analyzed using FlowJo software version 10.8.1 for Mac OS X. Phagocytic Scores were determined as (%*Mtb*^+^Live Cells x MFI of *Mtb*^+^ Cells)/100 for each of the antibody conditions tested.

### Binding affinity determination for LAM mAbs

ELISA plates (ThermoFisher 269620 439454) were precoated with 100 μL PBS containing 2ug/mL LAM overnight at 4°C. Plates were washed 5 times with PBS Tween 0.5% (PBST), blocked with 5% BSA for one hour at RT, and was again washed PBST 5 times before adding diluted mAb. Glycolipid-specific mAbs (SMITB14, MoAb1, CS-35, A194, OM2-L22, and CP2-R3, Table 1) at a starting concentration of 200 ng/mL were serially diluted 2-fold, and 9 dilutions per antibody were added to LAM-coated and pre-washed plates. Antibodies were incubated at RT for 2 hours before washing the plates and adding hIgG1 detection antibody (Bethyl Laboratory #A80-104P). Plates were incubated for 1 hour with secondary antibody and washed with PBST 5 times before adding TMB substrate (BD) to develop the signal The reaction was stopped with 2N H2S04 and ELISA plates were read on a Tecan infinite M1000pro plate reader. The collected data of the antibody dilution series was plotted and IC-50 was determined using GraphPad Prism.

### Luminex based FcR-binding assay

LAM was covalently coupled to MagPlex Luminex beads as described ^42^ and these beads were combined with 10 ng of αLAM Fc-variants and incubated for 1hr at RT to form immune complexes that were then washed with PBS and probed for binding to mouse FcγR. To measure FcγR binding to these Fc-variant immune complexes, FcγR detectors were generated by biotinylating Avi-tagged mouse FcγRIIB, FcγRIIA, RcγRIV (generated by the Duke Human Vaccine Institute) and conjugating the biotinylated FcγR with Streptavidin-PE (Prozyme PJ31S)^63^. FcR detectors were incubated with αLAM ICs, then washed and resuspended in BioRad Sheath Fluid. The MFI of FcγR-PE was detected for each condition on the FlexMap3D (Luminex).

### Flow staining of infected lungs

Single cell suspensions from *Mtb*-infected lungs were generated by digestion with 10mg/mL Type IV collagenase D (Worthington) and 30 μg/mL DNAseI (Roche) for 1hr at 37°C in a shaker. Cells were washed with PBS then stained for innate immune cells using viability dye (eBioscience Fixable Viability Dye eFluor455UV) and the fluorescently-labelled Abs BUV737-CD19 (1D3, BD), BUV395-CD45 (30-F11, BD), PerCp-CD11c (N418,Bioegend), BUV805-CD11b (M1/70), BV605-Ly6G (HK1.4, Biolegend), AlexaFluor700- Ly6C (1A8, Biolegend), Pacific Blue-IA-IE (M5/114.15.2, Biolegend), BV510-CD24 (M1/69, Biolegend), PE-Cy7-CD64 (X54-5/7.1 FC, Biolegend), BV711-CD16/32 (93, Biolegend), BV786-CD16.2 (9e9, Biolegend), and PE-CD351(TX61, Biolegend). *Mtb*-infected cells were identified by the expression of YFP. Lung cells were stained, washed with PBS, and fixed in 4% PFA overnight. The samples were analyzed using BD TruCount (Cat# 340334) tubes in order to determine the total number of CD45+ cells in each sample; the cells and TruCount beads were collected on BD FACS Symphony and analyzed with FlowJo version 10.8.1 for Mac OS X. The gating strategy for immune cell identification is outlined in Fig. S4.

### Mtb restriction assay in sorted lung immune cells

Single cell suspensions were generated from lungs of naïve 6-week-old female C57Bl/6 mice and stained with antibodies as described in the methods used for flow cytometry of *Mtb*-infected lungs. AM, IM, and PMN were sorted from the lung suspension using the gating strategy outlined in Fig. S3 on a BD FACS AriaFusion. The cells were resuspended in phenol-free complete RPMI-10 media without antibiotic and plated in 96- well plates (Greiner BioOne: #655083) and 10,000 AM, IM, and PMN were plated per well. At least five days prior to infection of these cells, H37Rv-276, a luminescent protein-expressing *Mtb* (lux-*Mtb*), was cultured in 7H9-OADC media to mid-log phase at 37°C and passaged once prior to infection. Bacteria was washed and resuspended in RMPI-10 without phenol and pre-coated with antibody as described above for *Mtb* surface staining. 1.5 x10^4^ antibody-coated *Mtb* was added to 96-well plates containing previously aliquoted lung immune cells for an approximate MOI of 0.5. Luminescence readings were taken daily over the course of 192 hours. Luminescent values were fold normalized to background signal captured in uninfected wells of the plate, and background-corrected luminescent values were reported as the fold change in lux-*Mtb* signal relative to the starting *Mtb* luminescence at the 0 hr time point of the assay. Area Under the Curve (AUC) values for each mAb treatment *Mtb* growth curve was calculated to quantify the difference the bacterial restriction mediated by mAb treatment.

### Single-cell sorting and RNA Sequencing of mouse lung cells

6-week-old female C57Bl/6 mice were purchased from Jackson Laboratories and injected with PBS or 100 μg of aLAM-mIgG2a or -mIgG2a N297A (5 mg/kg) i.p. (3 mice/group) one day prior to aerosol infection (Biaera AeroMP- Hope aerosolization unit) with YFP-expressing H37Rv. On day 5 post infection, mice were sacrificed, and a single cell suspension of lungs cells was generated for flow sorting. Prior to sorting for CD45+ cells, samples were then quenched with FACS buffer (PBS with 2% FBS and 2 mM EDTA), washed once, and counted before proceeding with MULTI-seq barcoding. Samples were barcoded with 2.5 μM of the lipid-modified oligo (LMO) anchor and barcode for 5 minutes on ice in PBS before adding 2.5 μM of the LMO co-anchor and incubating for an additional 5 minutes. Samples were quenched with 1% BSA in PBS and washed once with PBS before staining with the antibodies according to the above flow staining method. CD45+ cells were sorted on a BD FACS AriaFusion. Sorted cells were processed using the 10X Genomics NextGEM Single Cell 3’ kit v3.1 per the manufacturer’s protocol in 2 microfluidic lanes. Again, 0.5 U/μL RNase inhibitor (Roche) was added to the single-cell suspension and cDNA was heat-inactivated at 95°C for 15 minutes prior to BSL3 removal. Libraries were sequenced on a NextSeq500 (Illumina), and the data were aligned to the mm10 reference using Cell Ranger Count v6.0.1.

### Analysis of infection scRNA-seq data

LMO barcode and gene expression count matrices were merged and analyzed using R (v4.0.3) and Seurat (v4.0.0). Samples were demuxed using HTODemux (Seurat). Data from both CD45+ and *Mtb*+ sorted samples were merged. Cells with less than 200 unique molecular identifiers (UMIs) counted and more than 25% mitochondrial UMIs were excluded. 3,000 variable features were used for PCA. Counts were normalized using the default parameters from NormalizeData (Seurat), i.e., scaling by 10,000 and log normalization. Walktrap (igraph) clustering was performed on the shared nearest neighbor graph generated from FindNeighbors (Seurat) using 20 principal components and k = 20. Cell type annotation was based on expert annotation and predicted cell type labels from the Tabula Muris dataset. Cell type labels were predicted using FindTransferAnchors, MappingScore, and TransferData (Seurat) with 20 dimensions and 20 trees. Myeloid cell types were subclustered separately by repeating the steps above on the cell subsets. Marker gene statistics were calculated using wilcoxauc (presto). The *in vivo* scRNA-seq data is accessible on GEO (GSE198064).

### Gene Set Enrichment Analysis

Pathway enrichment analysis was performed comparing PBS-, mIgG2a-, and mIgG2a N297A-treated mice to the others (mIgG2a/PBS, mIgG2a/N297A, and N297A/PBS). GSEA software (version 4.2.1) was used to rank the genes expressed in AM from treated mice and calculate enrichment of genes in hallmark genes list curated by MSigDB^64^. Gene-sets enriched in either treatment group with a nominal p value < 0.05 were used for bubble plot visualization.

### Data Analysis and Visualization

Univariate data visualization and statistical analysis were performed using GraphPad Prism (Version 9.3.1) and “ggplot” package in RStudio (version 2021.09.0+351).

## Data availability

In addition to data included within the main text and supplemental materials, Flow cytometry and meta data associated with mouse immune profiling can be found online (https://fairdomhub.org/studies/927) along with the tabulated immune cell measures derived from flow files and gates outlined in Fig. S4. scRNAseq files, meta data, and analysis code can be accessed no GEO (GSE198064).

## Materials availability

Monoclonal antibodies generated in this study will be made available on request with a completed materials transfer agreement.

## Acknowledgements

We thank the Dana Farber Cancer Institute for large scale antibody production. We would like to thank the Ragon Institute Flow Cytometry for expertise and technical assistance with sample analysis. In addition, we are grateful the BioMIcro Center, Data Management and Analysis Core of MIT for data management support. We thank the Duke Human Vaccine Institute for providing us with the avi-tagged mouse FcR reagents.

## Funding

Ragon Institute of MGH, MIT, and Harvard and the SAMANA Kay MGH Research Scholar Program (GA), Bill and Melinda Gates Foundation grant OPP1156795 (GA and SMF). National Institutes of Health definitive contract no.75N93019C00071 (SMF, GA, BDB, CMS). National Institutes of Health grant T32 AI007061 (PSG). Yerby Fellowship Program (PSG).

## Author Contributions

Conceptualization: PSG, GA, SF. Methodology: PSG, JMP, JS,RL, BAF, AV, MDS, EBI, MKP, CMS, AC, AP, RR, C. Locht, GK, RDC, MHW, TM, BB, GA, SF. Investigation: PSG, GA, SF. Visualization: PSG, JMP. Funding acquisition: GA, SF. Project administration: C. Luedeman Supervision: GA, SF. Writing – original draft: PSG, GA, SF. Writing, review, & editing: all authors.

## Competing interests

GA is a founder of SeromYx Systems and is an employee of Moderna. All other authors have no conflicts to declare.

## Supplementary Figure Legends

**Supplementary Figure 1: Protein antigen biding profiles of novel monoclonal antibodies.** The novel antibody clones derived from vaccination against capsular (CP) and outer membrane (OM) fractions of *Mtb* cell surface were screened for binding of *Mtb* protein antigens in a customized Luminex bead assay. Graphed are the antigens which displayed binding signal above isotype control: **(A)** Esat6 and CFP10 proteins, **(B)** Psts1, and **(C)** HspX.

**Supplementary Figure 2: Antibody mediated phagocytosis of *Mtb* and the relationship with surface binding and *in vivo* control.** Monoclonal antibodies were combined with yellow fluorescent protein expressing *Mtb* (YFP-*Mtb*) for 1 hour to form immune complexes before combining with bone marrow-derived dendritic cells, neutrophils, and macrophages. (**A**) Gating strategy to identify and quantify the frequency of YFP-Mtb+ cells and the MFI of the YFP signal within infected cells, and phagocytosis (Phago) Scores were determined using these values. Phago Score = ((%*Mtb*^+^cells x *Mtb*^+^Cell mean fluorescence intensity)/100). (**B**) Pearson correlation analysis found moderate to weak linear relationships between of in vivo *Mtb* growth restriction (*Mtb* fold change in vivo), *Mtb* surface binding, DC phago, neutrophil phago, and macrophage phago. Pearson r summarized the correlation between measures.

**Supplementary Figure 3: Gating strategy to identify immune cell subsets in the lungs of Mtb infected mice**. Representative gates outline the strategy used to identity populations of alveolar macrophages (AM), eosinophils, neutrophils (PMN), B cells, CD103+ DC, T cells/NK cells, Ly6C^high^ Monocytes, Ly6C^low^ Monocytes, recruited macrophages and CD11b+ DC.

**Supplementary Figure 4:** α**LAM Fc-variants do not promote *in vitro* restriction of *Mtb*.** Growth curve of luminescent *Mtb* (lux-*Mtb*) in **(A)** alveolar macrophages, **(B)** interstitial macrophages, and **(C)** neutrophils sorted from naive mouse lung tissue were infected with *Mtb* pre-opsonized with aLAM-mIgG2a, -mIgG2a N297A variants or isotype control mAbs. x,y plots depict the fold-increase over day 1 luminescence. Bar plots depict area under the curve (AUC) of luminescence measured in the x,y, plot over 8 days. Graphs depict the mean +/- SEM. No significant differences between Fc-variants were identified for AUC in one-way ANOVA with Tukey’s multiple correction.

**Supplementary Figure 5: Bacterial Burden and analysis of immune cells present at Day 5 post infection.** C57BL/6 mice were pre-treated with PBS or 100mg of mAb mIgG2a or mIgG2a N297A aLAM mAb and infected via aerosol with ∼60 Day 1 CFU of YFP-*Mtb.* **(A)** Similar levels CFUs present in the lungs treated lungs 5 days after aerosol infection with H37Rv-YFP. **(B)** Relative frequency (%) of cell types identified by scRNAseq analysis. **(C)** Relative % of immune cells identified by flow cytometric analysis (as defined in supplemental figure 4).

**Supplementary Figure 6: FcR expression on immune cells recruited to the lung following *Mtb* infection.** Relative expression of activating receptors (FcgRI, FcgRIIA, FcgRIV) and inhibitory FcgRIIB on innate immune cell populations recruited to infected lungs at 14 days post infection with ∼50 Mtb CFU. Immune cells identified by flow cytometric analysis (as defined in supplemental figure 4).

**Supplementary Table 1:** Screening of *Mtb* outer membrane and capsule antibody clones from vaccination. Antibodies were probed for binding to *Mtb* antigens via western blot, microscopy, and whole *Mtb* ELISA. *Western blots* performed on outer membrane preparations (OM2), capsule preparation (C1), and purified protein (SOD1), band size associated with antibody binding are listed in the above corresponding blotted *Mtb* fractions. *Mtb ELISA Binding* is defined as “Specific” for *Mtb*, “Weakly Specific” based on low binding to *Mtb* or some binding to irrelevant bacteria (*E.coli*), “Non-specific” for *Mtb* based on lack of binding or significant binding to irrelevant bacteria (*E.coli*). Finally, ELISA was performed on *Mtb* cells grown in the presence or absence of Tween-80, which removes peripherally-associated cell envelope antigens, and the ratio of signal with and without detergent was determined. Low Detergent: No Detergent binding ratios suggest mAb clones associate with Mtb-envelope associated antigen.

| clone name | Western Blot: OM2 (KDa) | Western Blot: C1(KDa) | Western Blot: SOD1(KDa) | Mtb ELISA Binding* | Detergent: No Detergent ELISA Signal Ratio |
| --- | --- | --- | --- | --- | --- |
| C1-LR/NP-12 | 32.3 |  | 46.1 | Non-Specific | 0.78 |
| CP2-LR17 | 24.3, 15.4 | 15.4 |  | Specific | 0.52 |
| CP2-LR5 | 41.1 | 41.1 |  | Non-Specific | 0.54 |
| CP2-R19 |  |  |  | Non-Specific |  |
| CP2-R3 |  |  |  | Specific | 0.28 |
| OM2-L2 | 15.2 | 15.2 |  | Specific | 2.84 |
| OM2-L22 |  |  |  | Weakly Specific | 1.80 |
| OM2-L7 |  |  |  | Specific | 0.61 |
| OM2-R18 |  |  |  | Non-Specific |  |

## Notes

### Competing Interest Statement

The authors have declared no competing interest.

