## Supplemental Figures for "Antibody-Fab and -Fc features promote *Mycobacterium tuberculosis* restriction"

### Supplemental Figure 1

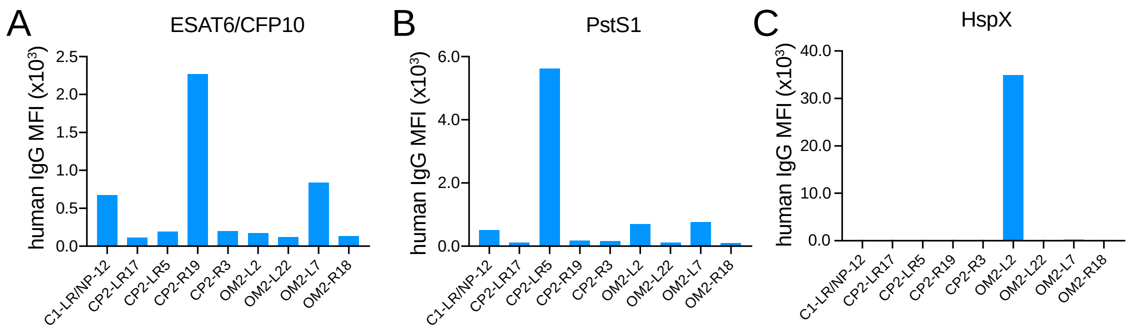

### Supplemental Figure 2

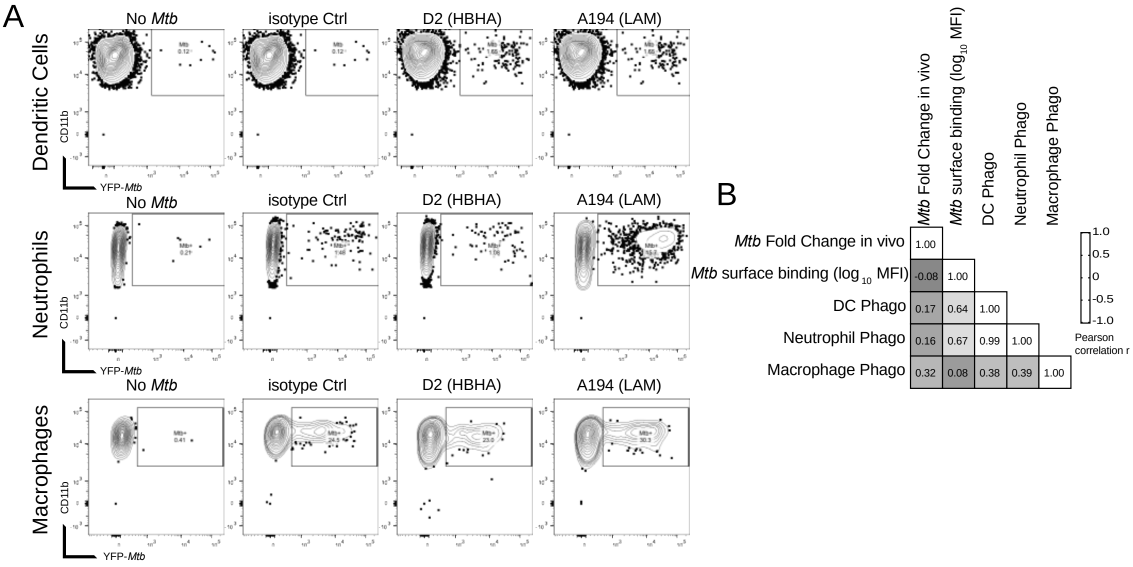

Supplemental Figure 3

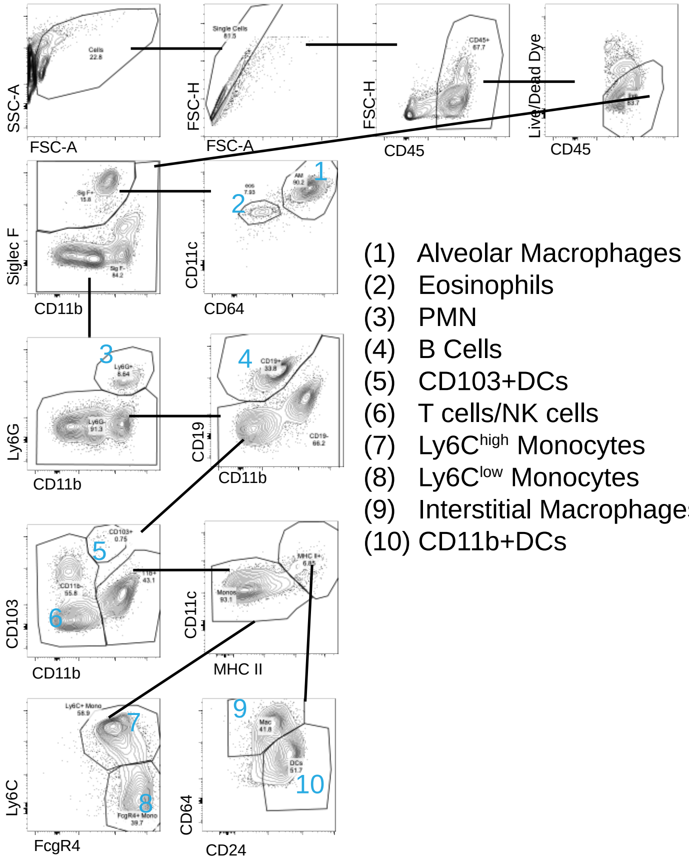


(1) Alveolar Macrophages
(2) Eosinophils
(3) PMN
(4) B Cells
(5) CD103+DCs
(6) T cells/NK cells
(7) Ly6C$^{high}$ Monocytes
(8) Ly6C$^{low}$ Monocytes
(9) Interstitial Macrophages
(10) CD11b+DCs

### Supplemental Figure 4

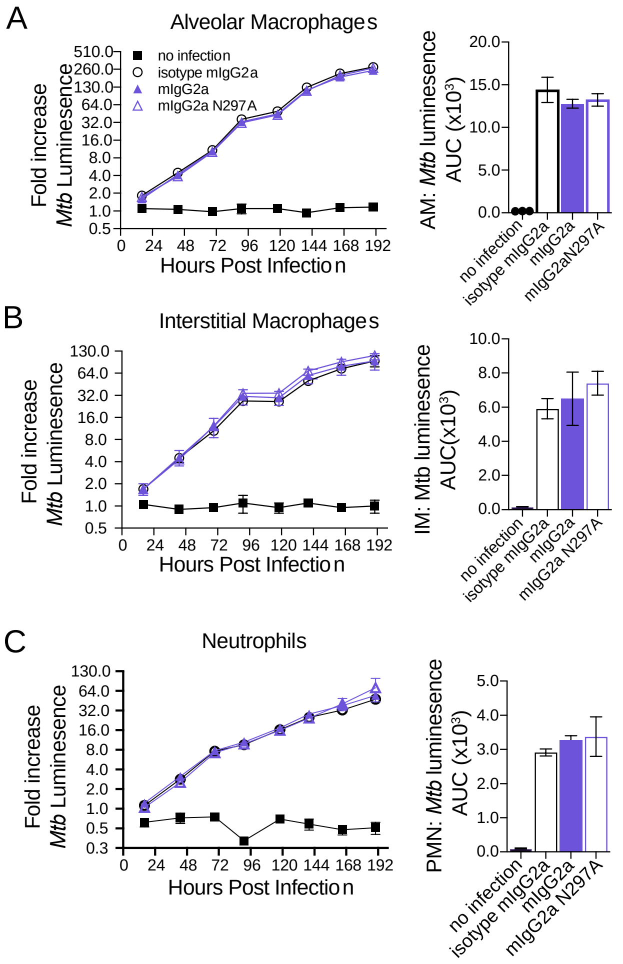

Supplemental Figure 5

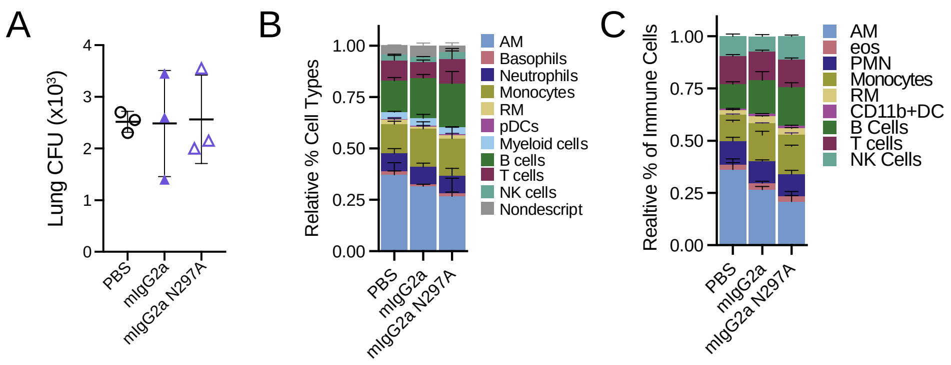

### Supplemental Figure 6

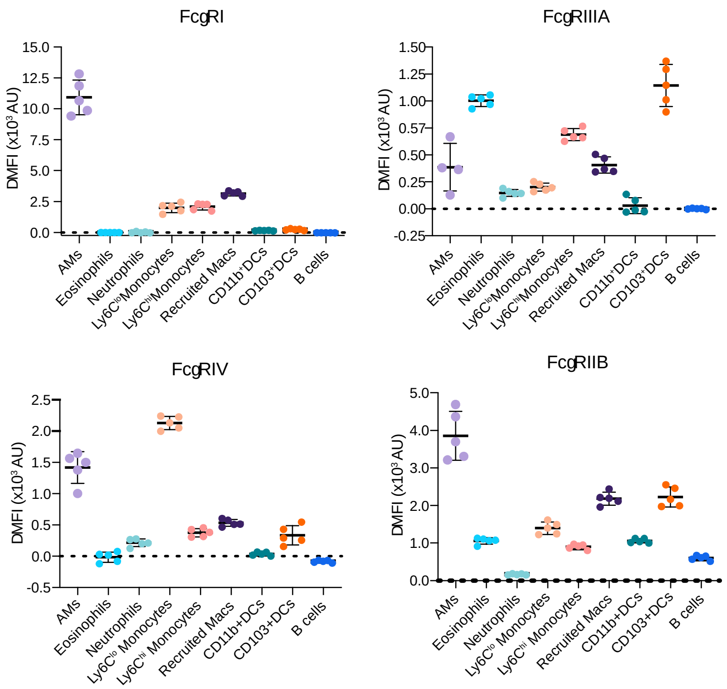
